# Chronic IFNα treatment induces leukopoiesis, increased plasma succinate and immune cell metabolic rewiring

**DOI:** 10.1101/2022.10.17.512487

**Authors:** Anjali S. Yennemadi, Gráinne Jameson, Mary Glass, Carolina De Pasquale, Joseph Keane, Massimiliano Bianchi, Gina Leisching

**Affiliations:** TB Immunology Group, Department of Clinical Medicine, Trinity Translational Medicine Institute, St James’s Hospital, Trinity College Dublin, The University of Dublin, Dublin, Ireland; Ulysses Neuroscience Limited, Trinity College Institute of Neuroscience, Lloyd Institute, Trinity College Dublin, Ireland

**Keywords:** interferon alpha, interferon therapy, glycolysis, oxidative phosphorylation, metabolism, mitochondria, macrophages

## Abstract

Although clinically effective, the actions of IFNα, either produced endogenously or by therapeutic delivery, remain poorly understood. Emblematic of this research gap is the disparate array of notable side effects that occur in susceptible individuals, such as neuropsychiatric consequences, autoimmune phenomena, and infectious complications. We hypothesised that these complications are driven at least in part by dysregulated cellular metabolism. Male Wistar rats were treated with either 170,000 IU/kg human recombinant IFNα-2a or BSA/saline (0.9% NaCl) three times per week for three weeks. Bone marrow (BM) immune cells were isolated from the excised femurs for glycolytic rate and mitochondrial function assessment using Agilent Seahorse Technology. Frequencies of immune cell populations were assessed by flow cytometry to determine whether leukopoietic changes had occurred in both blood and BM. Plasma levels of lactate and succinate were also determined. BMDMs were metabolically assessed as above, as well as their metabolic response to an antigenic stimulus (iH37Rv). We observed that BM immune cells from IFN-treated rats exhibit a hypermetabolic state (increased basal OCR/GlycoPER) with decreased mitochondrial metabolic respiration and increased non-mitochondrial OCR. Flow cytometry results indicated an increase in immature granulocytes (RP1-SSChi CD45lo) and classical monocytes (CD43lo RP1hi) in the blood, together with increased succinate levels in the plasma. BMDMs from IFN-treated rats retained the hypermetabolic phenotype after differentiation and failed to induce a step-up in glycolysis and mitochondrial respiration after bacterial stimulation. This work provides the first evidence of the effects of IFNα treatment in inducing hypermetabolic immune features that are associated with markers of inflammation, leukopoiesis, and defective responses to bacterial stimulation.

## 1. Introduction

Interferon (IFN)α therapy has been used extensively for its efficacy in the treatment of various disease conditions including chronic hepatitis, lymphoproliferative disease, and cancer. However, IFNα treatment is associated with a wide variety of side effects that often outweigh the clinical benefit [1]. These include cognitive impairments and neuropsychiatric consequences, and in some cases, the development of autoimmune phenomena [2] and infectious complications [3]. Thus, although the immunomodulatory effects of IFNα in the treatment of disease have benefits, there also lies the potential for pathogenesis[4].

Chronic exposure to IFNα during therapy has been shown to induce immune cell activation which is hypothesised to drive the development of these autoimmune conditions [5] however, the mechanisms underpinning these findings are unclear. Research has shown that dysregulated metabolism is associated with immune cell dysfunction and this has implications for inappropriate immune activation and chronic inflammation[5]. Therefore, we hypothesise that IFNα treatment induces metabolic reprogramming that may contribute to IFNα-induced pathogenesis.

In this study, using *in situ* methods, we provide a deeper understanding of IFNα-induced metabolic changes that occur at the level of the bone marrow, as well as well as the blood, from an immunometabolic perspective.

## 2. Methods

### 2.1 Animals

Male Wistar rats (8/10 weeks old) sourced from Envigo UK, were pair-housed in a controlled environment (temperature: 20–22°C, 12/12 hr light/dark cycle (lights on at 8am), with water and food ad libitum. Upon arrival, animals were randomly paired into cages. The first 2 animals of every block were assigned to the saline group, allowing the saline cages to be placed randomly in the stack. Animals were acclimatised to the facility environment for 2 weeks and handled by experimenters before starting experiments. Animal weight and food consumed was monitored throughout the experiment, no animals were excluded from the study. Treatments are alternated between animals so that no two animals are treated with the same drug directly after one another. Nine units per group were used, therefore 18 units in total. OCR, GlycoPER, immune cell frequencies, plasma lactate and succinate were outcome measures.All experiments were performed under license from the Health Products Regulatory Authority (HPRA) of Ireland following EU regulations and with local ethical approval (The University of Dublin, Trinity College Dublin).

#### 2.1.1 IFN-alpha treatment and Compound Administration

IFN-α treatment was administered as previously published[6]. Animals were randomly assigned to be treated with either 170,000 IU/kg human recombinant interferon alpha-2a (Merck), or BSA/saline (0.9% NaCl) three times per week for three weeks with n=9 per group. All injection solutions were prepared in 0.1% BSA/saline and administered at a volume of 0.3 ml/kg subcutaneously. All rats were anesthetised by CO_2_ inhalation and decapitated. Trunk blood and femurs from single rats were collected and processed immediately. Researchers conducting ex vivo work were blinded to treatment groups.

### 2.2 Cell culture and treatment

Total BM immune cells were isolated from femurs, red blood cells were lysed, and cells were counted and plated in Agilent Seahorse XF plates at 150 000 cells/well or 200 000 cells/well for BM-derived macrophages (BMDMs). BM immune cells were cultured in DMEM supplemented with 10% FBS. Monocytes from BM were differentiated into macrophages with the addition of 20 μg/ml M-CSF (Cat#556904, BioLegend) after 7 days of culturing. For antigenic stimulation experiments, a precalculated volume (equating to a multiplicity of infection of 1-5) of irradiated H37Rv (iRv) was added to port A of the Agilent Seahorse flux cartridge for induced assays.

### 2.3 Metabolic analysis

Mitochondrial function was assessed using the Mitostress test and glycolysis was assessed using the Glycolytic rate assay to obtain oxygen consumption rate (OCR) and glycolytic proton extrusion rate (GlycoPER) readouts respectively, using a Seahorse XF extracellular flux analyser (Seahorse Bioscience, Inc, North Billerica, MA, USA). Seahorse medium was supplemented with 4.5 g/L D-glucose, 2 mM glutamine and 100mM pyruvate and subsequently added to the cells. Following incubation in a CO2-free incubator at 37 °C for 60 min, basal OCR and ECAR were recorded for 30 min. The MitoStress assay was performed by sequential addition of 2 μg/ml oligomycin, 0.7 μM carbonyl cyanide 4-(trifluoromethoxy) phenylhydrazone and 1 μM rotenone/antimycin A. The glycolytic rate assay was performed through the sequential addition of 1 μM rotenone/antimycin A followed by 5mM of 2-DGOCR and GlycoPER values were normalised using the Crystal Violet dye extraction growth assay and the Wave Desktop 2.6 Software (https://www.agilent.com) was used for analysis.

### 2.4 Flow cytometry

Isolated immune cells from BM and blood were immunophenotyped after resuspension in 50uL of BD Horizon Brilliant Stain Buffer (BD Biosciences), and stained for 10 mins at RT to detect classical and non-classical monocytes, immature granulocytes, neutrophils, B cells, CD4+ Tcells, CD8+ T cells,and then fixed in 1% PFA for 20 mins (Table 1). Cells acquired on an Amnis CellStream (Luminex Corporation, Austin, TX, USA) and gated as previously described[7]. Data analysis was performed using FlowJo software (Version 10.8.1, FloJo LIC, Oregon, USA).

**Table 1.** Antibodies/stains used to detect immune cell subsets in blood and bone marrow from IFNα- and saline-treated rats

| Antibody | Fluorochrome | Manufacturer | Clone | Category Code |
| --- | --- | --- | --- | --- |
| Live-Dead | eFluor780 | eBioscience | N/A | 65-0865-14 |
| CD11b | V450 | BD Biosciences | WT.5 | 562108 |
| CD3 | VioGreen | Miltenyi Biotec | REA223 | 130-119-125 |
| RP1 | BV605 | BD Biosciences | RP-1 | 743055 |
| CD4 | BV711 | BD Biosciences | OX-35 | 740723 |
| CD8a | BUV496 | BD Biosciences | OX-8 | 741106 |
| CD90 | APC | eBioscience | HIS51 | 17-0900-82 |
| CD45R | FITC | eBioscience | HIS24 | 202205 |
| CD161 | BB700 | BD Biosciences | 10/78 | 746225 |
| CD43 | PE-Vio770 | Miltenyi Biotec | REA503 | 130-107-686 |
| IgM | PE | eBioscience | HIS40 | 12-0990-82 |
| CD45 | PE-Dazzle 594 | BioLegend | OX-1 | 202223 |
| CD32 (FcR block) | N/A | BD Biosciences | D34-485 | 550270 |

### 2.5 Plasma lactate and succinate measurements

Plasma levels of lactate (Cat#J5021, Promega) and succinate (Cat#MAK355-1KT, Sigma) were assessed according to the manufacturer’s instructions.

### 2.6 Statistical analysis

Statistical analyses were performed using GraphPad Prism 9 software. Statistically significant differences between two normally distributed groups were determined using Student’s paired t-tests with two-tailed P-values. Differences between three or more groups were determined by one-way ANOVA with Tukey’s multiple comparisons tests. *P*-values of <0.05 were considered statistically significant.

## 3. Results

### 3.1 IFNα treatment induces metabolic dysregulation in BM cells and increased succinate concentrations and frequencies of immature granulocytes and classical monocytes in blood

We first sought to determine the overall metabolic phenotype of immune cells derived from the BM of IFNα-treated rats since the BM is a reservoir for long-lived plasma cells that infiltrate tissue. Basal GlycoPER levels were significantly higher in IFNα-treated rats (Fig.1A) and the mitochondria of these cells had a lower maximal respiratory capacity with a higher basal OCR (Fig. 1A). Non-mitochondrial oxygen consumption was also significantly increased in BM cells from IFNα-treated rats. We then investigated whether any leucopoietic changes in the bone marrow, and consequently, in the blood, had occurred. After flow cytometric assessment of B cell, T cell, monocyte and neutrophil popultaions, no change in cell frequencies were evident in the BM, however, in the blood of IFNα-treated rats we observed increased frequencies of immature granulocytes (RP1^-^, SSC^hl^ CD45^lo^) and total monocytes (CD43^lo^ RP1^hi^, and CD43^hi^ RP1^hi^), of which the classical monocyte subset (CD43^lo^ RP1^hi^) was dominant (Fig.1D). We also determined whether these metabolic pertubations affected circulating metabolites associated with glycolysis and inflammation, and found that increased concentrations of succinate, but not lactate, was evident in the plasma of IFNα-treated rats (Fig.1E).

**Figure 1.**
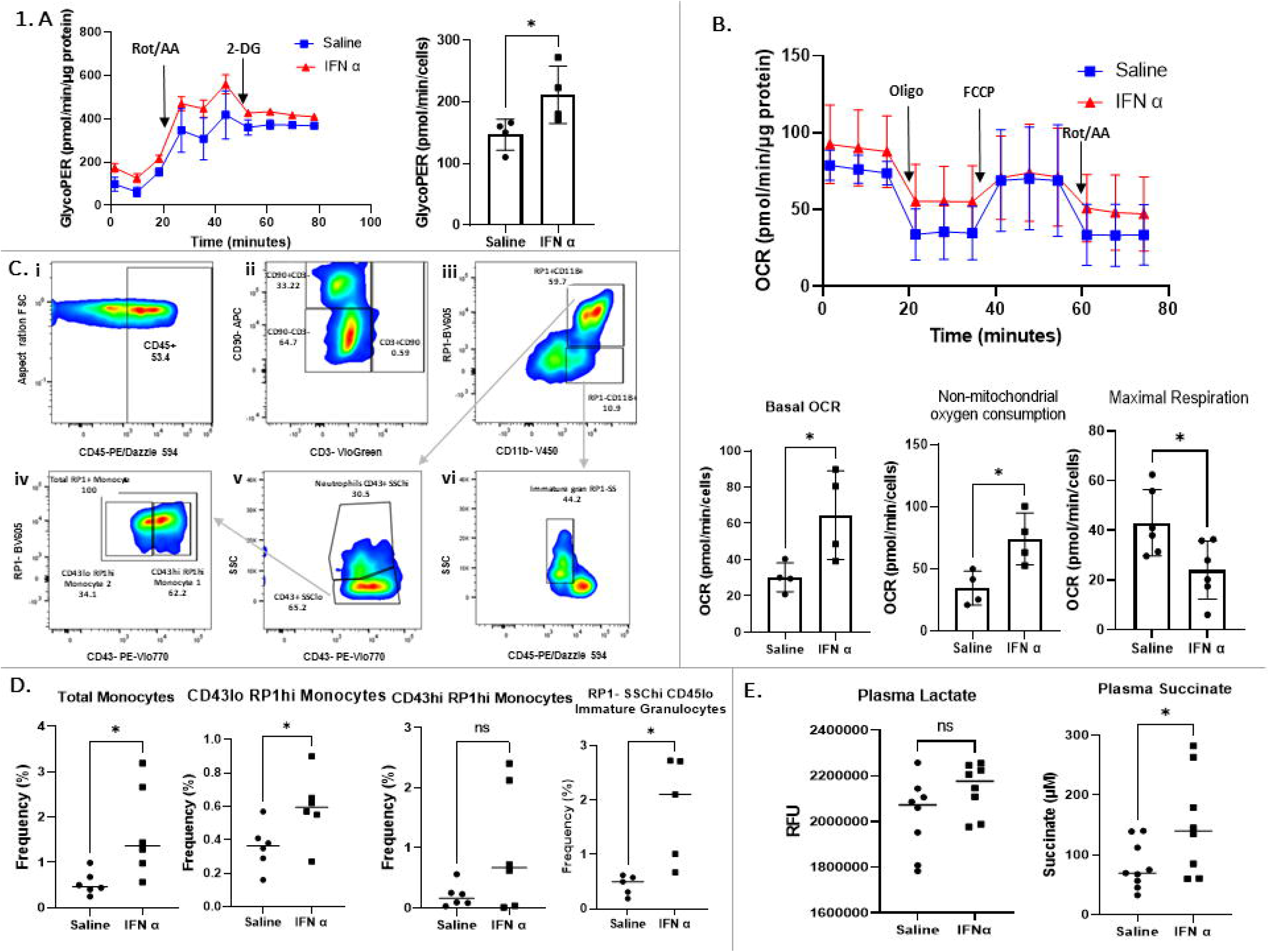
IFNα treatment induces metabolic rewiring and leukopoiesis of the BM as well as increased succinate levels in the plasma. **A**. The Glycolytic rate assay and Mitostress test revealed that immune cells from the BM of IFN-treated rats exhibited increased basal GlycoPER (n=4) and **B**. OCR with decreased maximal respiration and increased non-mitochondrial oxygen consumption vs. Saline treated rats (n=4). **C**. Flow cytometry gating strategy, clockwise: (C,i) Debris, doublets, and dead cells were excluded (data not shown) and CD45^+^ leukocytes gated (C,ii) CD90^−^CD3^−^ cells were agted on to exclude T and B cells. (C,iii) CD90^−^CD3^−^ was split into RP1^+^CDllb^+^ cells and RPl^−^CDllb+ cells. (C,vi) RPl^+^ cells were further gated into immature granulocytes by SSC^hi^ (RP^−^ CDllb^+^SSC^hi^). (C,v) RPl^+^CDllb^+^ cells were divided into neutrophils (CD43^+^SSC^hi^) and CD43^+^SSĆ° cells. (C,vi) SS^lo^ cells were divided into two monocyte subsets, CD43^lo^RPl^hi^ and CD43^hi^RPl^lo^· D. Flow cytometry analysis of the blood showed that IFN treated rats had increased frequencies of total monocytes, with CD43^lo^ RPl^hi^ monocytes accounting for the majority of the monocytes detected, as well as an increase in RPl-SSC^hi^ CD45^lo^ immature granulocytes (n=5). E. Assessment of plasma showed that only succinate was significantly elevated in the plasma of IFN treated rats vs. saline treated rats. Paired t test, *p<0.05.

### 3.2 BMDMs retain a dysregulated metabolic phenotype after culture and do not metabolically respond to a bacterial stimulus

Next we investigated if this dysregulated metabolic phenotype was retained after BM-derived monocytes were differentiated into macrophages. We observed that even after 7 days of culture and differentiation, both basal glycolysis (Fig.2A) and basal OCR (Fig.2B) were significantly increased in macrophages from IFNα-treated rats. Maximal respiratory capacity of mitochondria from IFNα-treated rats was significantly higher (Fig.2B), indicating metabolic rewiring that affected the functional capacity of these mitochondria in comparison to macrophages from saline-treated rats. We then determined whether this rewired macrophage metabolic response could provide insight into a possible mechanism underpinning infectious complications after IFNα treatment [3]. BMDMs from IFNα-treated rats stimulated with iRv failed to induce glycolysis and maximal respiration when compared to BMDMs from saline treated rats (Fig.2C). Their overall metabolic phenotype was blunted in comparison to the induced metabolic phenotype observed in the saline-stimulated BMDMs (Fig.2D).

**Figure 2.**
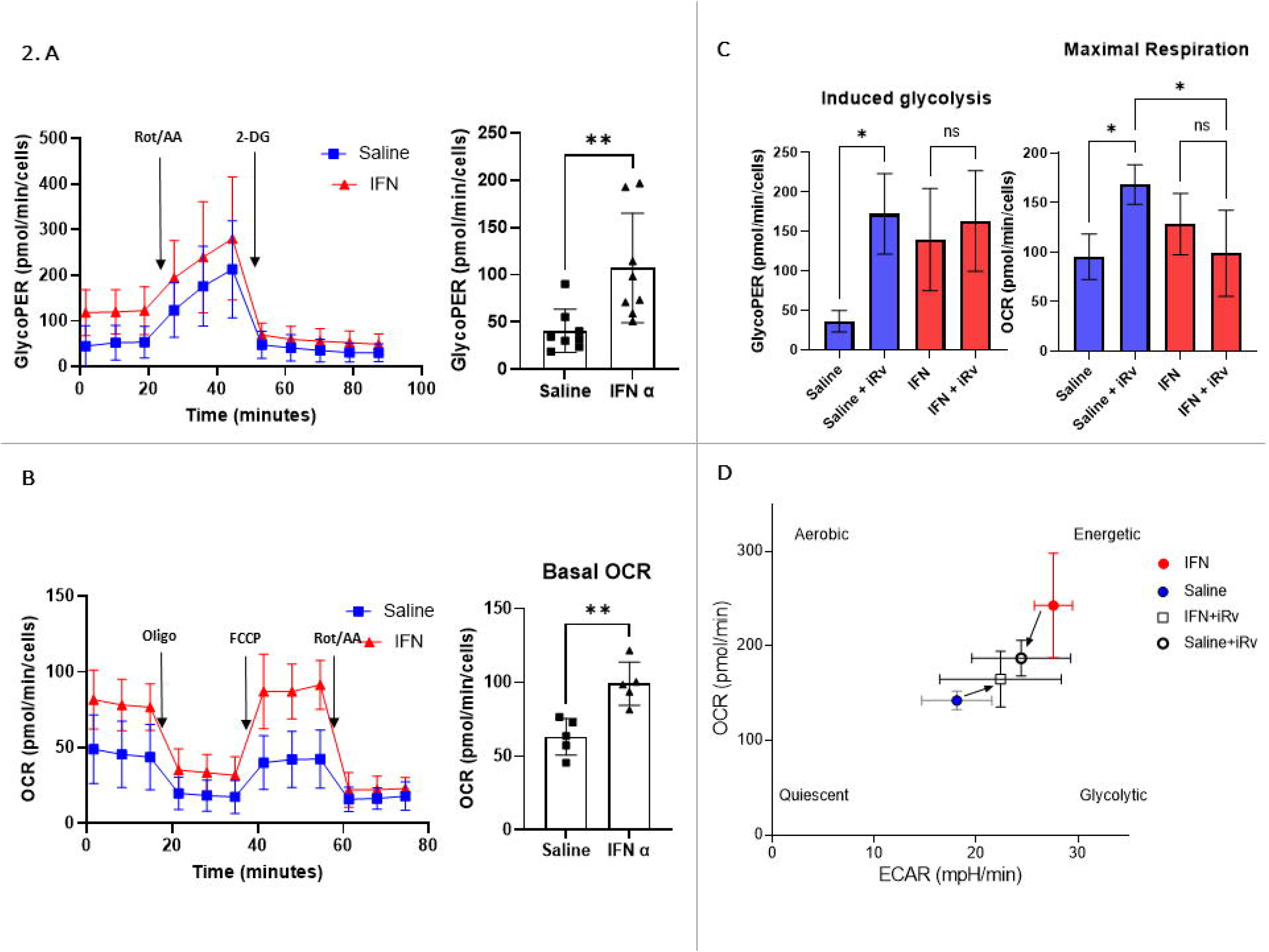
BMDMs from IFNα-treated rats are hypermetabolic and fail to metabolically respond to iH37Rv stimulation. **A**. BMDMs from IFN-treated rats exhibited increased basal glycoPER (n=8) and **B**. OCR (n=5) in comparison to saline-treated controls. **C**. BMDMs from saline-treated rats stimulated with iH37Rv (MOI 1-5) were able to significantly induce glycolysis and engage in maximal respiration vs. BMDMs from IFN-treated rats (n=3). D. Overall energetic phenogram depicting BMDMs before and after stimulation with iH37Rv indicating an abnormal energetic response of BMDM from IFN-treated rats towards quiescence (directed by arrows). Paired t test or one way ANOVA with Tukey’s multiple comparisons tests *p<0,05, **p<0,0l.

## 4. Discussion

This work sought to address the knowledge gap of the metabolic parameters which may contribute to IFNα therapy-associated immune cell activation and susceptibility to infection. Here, we suggest that dysregulated immune cell metabolism could at least, in part, support a mechanistic basis of these pathogenic side-effects that are observed in susceptible individuals.

We observed that BM immune cells are metabolically primed (Fig1A and B) with increased basal oxidative phosphorylation and glycolysis, suggesting a hypermetabolic state[8]. Notably, we observed an increase in non-mitochondrial OCR in BM immune cells from IFNα-treated rats, which is typically attributed to enzymes associated with inflammation and is a negative indicator of bioenergetic health [9]. Global changes in metabolism that occur in activated immune cells such as these often correlate with the presence of individual metabolites, including succinate, citrate, and NAD^+^ [10]. We observed increased succinate levels in the plasma of IFNα-treated rats (Fig.1E) which was not surprising as it has been described as a signal for inflammation and can accumulate in immune cells during inflammatory episodes [10, 11]. Elevated levels of circulating succinate is also evident in obese and diabetic rats [12], suggesting that in this model, increased succinate levels may indicate IFNα-induced changes in metabolism. In IFNα-treated rats, the presence of immature granulocytes in peripheral blood is indicative of leukopoiesis [13] and the increased frequency of classical monocytes may represent the earliest indicator of bone marrow stimulation by inflammation (Fig. 1D) [14].

A greater inflammatory burden is linked with a greater demand for ATP and higher metabolic activity, a feature that is commonly associated with inflammatory M1 macrophages[15]. We found that monocytes from IFNα-treated rats were pre-programmed to differentiate into macrophages that retained the metabolic phenotype observed in the BM. We were then interested in finding out whether these metabolically primed macrophages were able to appropriately respond to an antigenic stimulus in this metabolically heightened state. Interestingly, the BMDMs from IFNα-treated rats could not induce an appropriate metabolic response after bacterial stimulation. We propose that this may be due to their mitochondria having already reached maximal capacity, therefore rendering them unable to further engage OXPHOS in response to iH37Rv. This finding is in agreement with other work showing that type I IFN decelerate macrophage metabolism after infection with H37Rv [16]. This defective response is also in agreement with previous work that shows that normal mitochondrial respiration restricts growth of the bacterial pathogen *Listeria monocytogenes* [17] and that patients with mitochondrial disease have a higher susceptibility to infection [18]. It is therefore likely that other tissue resident cells, or those egressing from the BM, display features of dysregulated metabolism. This was observed in a longitudinal study that assessed whole-brain metabolic activity in patients by ^18^F-labeled fluorodeoxyglucose uptake before, and 4 weeks after, IFNα administration [19]. Widespread bilateral increases in glucose metabolism were observed in the basal ganglia and cerebellum, which correlated with self-reported fatigue. This supports the notion that IFNα-induced metabolic changes are systemic when IFNα supersedes its protective potential in susceptible individuals during therapy.

This is the first link between IFNα treatment and metabolic rewiring associated with with plasma markers of inflammation, leukopoiesis, and defective metabolic responses to bacterial stimulation. Metabolic interventions may therefore be a promising avenue of treatment for susceptible individuals receiving IFNα therapy.

## Abbreviation List

OCR: oxygen consumption rate
GlycoPER: Glycolytic proton extrusion rate
IFN: interferon
BMDM: bone marrow derived macrophages
M-CSF: macrophage colony-stimulating factor
BM: bone marrow
NAD^+^: Nicotinamide Adenine Dinucleotide

## 5. Author contributions

Conceptualisation; GL, MB, JK Data curation; GL, GJ,AY Formal analysis; GL,GJ Funding acquisition; JK, MB Investigation; GL, AY Methodology; AY, GJ,CDP, MG,GL, Resources; MB,JK Supervision; GL,JK,MB; Writing - original draft;AY,GL Writing - review & editing; AY,GL,GJ,JK,MB,MG

